# Mouse Classical and Non-Classical Monocytes Express Comparable Levels of Chemokine Receptor CX3CR1

**DOI:** 10.1101/2020.04.10.035550

**Authors:** Luca Cassetta, Esther Heideveld, Gabriele Matilionyte, Stamatina Fragkogianni, Emile van der Akker, Nicolle Kippen, Takanori Kitamura, Jeffrey W. Pollard

**Author notes:** Correspondence: Takanori Kitamura, 47 Little France, Edinburgh, Jeffrey W. Pollard, 47 Little France, Edinburgh, lead author. These authors contributed equally (Luca Cassetta, Takanori Kitamura).

## Abstract

Human and mouse monocytes are divided into two subpopulations, classical (C-Mo) and non-classical (NC-Mo) monocytes. CC-chemokine receptor 2 (CCR2) and CX3C-chemokine receptor 1 (CX3CR1) are common features of monocyte subsets between humans and mice, i.e., C-Mo and NC-Mo are characterized as CCR2^high^CX3CR1^low^ and CCR2^low^CX3CR1^high^. Since many studies utilize mouse models to investigate roles of monocytes in human diseases, it is important to understand the similarities and differences between human and mouse monocytic subsets. In this study, we show that the expression of *Cx3cr1* mRNA and CX3CR1 cell surface protein are different between circulating monocytic subsets in human but not in mice. We analyzed monocyte subsets in the blood using wild type C57BL/6 and *Cx3cr1*-GFP knock-in (*Cx3cr1*^GFP/+^) reporter mice. We observed higher *Cx3cr1* promoter activity indicated by GFP expression in NC-Mo compared to C-Mo. However, there were no differences between the subsets in CX3CR1 mRNA nor surface protein expression determined by anti-CX3CR1 antibody or binding of fluorophore-conjugated ligand. However in the bone marrow of *Cx3cr1*^GFP/+^ mice, CX3CR1 expression was higher in NC-Mo compared to C-Mo, suggesting that mouse NC-Mo express higher level of CX3CR1 than C-Mo in the bone but this difference disappears in the blood. In contrast, human NC-Mo differentially expressed CX3CR1 compared to C-Mo in both blood and bone marrow. Given these findings, the discrepancy between promoter activity and protein levels should be considered when the roles of CX3CR1 are investigated in mouse models of human diseases.

**Summary sentence:** In this study, we show that the expression of *Cx3cr1* mRNA is different between circulating monocytic subsets in human but not in mice.

## INTRODUCTION

Monocytes are innate immune cells that originate from the bone marrow and circulate in the blood. The circulating monocytes consist of at least two subpopulations, i.e., classical monocytes (C-Mo; CD14^+^CD16^-^ in human, Ly6C^+^ in mouse) and non-classical monocytes (NC-Mo; CD14^dim^CD16^+^ in human, Ly6C^-^ in mouse), which possess distinct biological functions [1, 2]. For example, C-Mo (also known as ‘inflammatory’ monocytes) migrate into inflamed tissue and differentiate to macrophages and dendritic cells, where they participate in the resolution of inflammation, facilitate wound repair, and enhance bacterial clearance [3]. C-Mo are also recruited into tumors and differentiate into a distinct population of macrophages, where they promote tumor progression [4]. In contrast, NC-Mo (also called ‘patrolling’ monocytes) survey the blood vessel and contribute to vascular maintenance, an early immune response, and tumor rejection [5-7]. As such, monocyte subsets therefore play distinct roles in disease progression and consequently they may represent important therapeutic targets. To understand the roles of monocyte subsets in human diseases, mouse models are essential and thus similarities between human and mouse monocyte subsets have been investigated [8]. One of the important similarities between human and mouse monocyte subsets is a distinct expression profile of chemokine receptors, i.e., CC-chemokine receptor 2 (CCR2) on C-Mo and CX3C-chemokine receptor 1 (CX3CR; also known as the fractalkine receptor) on NC-Mo, respectively in both species. [8-10]. In order to define CX3CR1 levels, most studies in mice utilize a *Cx3cr1*-GFP reporter model in which the *Cx3cr1* gene is replaced by a green fluorescent protein (GFP) gene that is transcribed from the *Cx3cr1* promoter [11]. In this model, NC-Mo compared to C-Mo express higher levels of GFP suggesting higher expression of CX3CR1 [10]. Based on these findings, CCR2 and CX3CR1 have been used as useful markers to distinguish C-Mo (CCR2^high^CX3CR1^low^) and NC-Mo (CCR2^low^CX3CR1^high^) both in human and mouse. However, GFP intensities in the *Cx3cr1*-GFP knock-in reporter mice do not necessarily reflect actual CX3CR1 protein levels, and thus differences in CX3CR1 mRNA or protein levels between mouse NC-Mo and C-Mo have not been explored. In this study, we report that circulating mouse C-Mo and NC-Mo express comparable levels of CX3CR1 mRNA and protein whereas human NC-Mo express higher levels of CX3CR1 than C-Mo. In contrast to monocytes in the blood, mouse as well as human bone marrow NC-Mo expressed significantly higher levels of CX3CR1 compared to C-Mo (**Supplementary Figure 1**; graphical abstract).

## MATERIAL AND METHODS

### Human Blood and Bone Marrow Samples

All study protocols were approved by the University of Edinburgh Ethics Committee (Edinburgh, UK) and the Medical Ethical Review Board of the Amsterdam Medical Center (Amsterdam, The Netherlands). Informed consent was obtained from all donors. Blood samples (10-100 mL) were collected from healthy female donors through the CIR Blood Resource (AMREC: #15-HV-013) and transferred into EDTA coated tubes (S-Monovet, Sarstedt). Human adult bone marrow aspirates (3-10 mL) were collected from the sternum of patients undergoing cardiac surgery in the Amsterdam Medical Center (MEC:04/042#04.17.370), and transferred to heparin coated tubes (Greiner). All samples were immediately stored at 4 °C and processed within 1 hour.

### Mouse Blood and Bone Marrow Samples

All procedures involving mice were conducted in accordance with licensed permission under the UK Animal Scientific Procedures Act (1986) (Home Office license number P526C60B3). Wild type mice on a C57BL/6 background were obtained from Charles River, and *Cx3cr1*^GFP/+^ and *Cx3cr1*^GFP/GFP^ mice [11] were kindly donated by Dr. Judith Allen, University of Manchester. Animals were housed and bred under standard conditions of care. Up to 500 μL of blood was collected from the heart of mouse (female, 8-12 weeks old), and transferred to tubes including heparin solution (5 IU/tube). The bone marrow cells were isolated from the tibias and femurs by flushing the bone with phosphate-buffered saline (PBS; Invitrogen). Samples were stored on ice until use.

### Quantitative PCR Analysis

Total RNA was extracted from sorted monocytes using the RNeasy Microkit (Qiagen) according to manufacturer’s instructions. Total RNA (0.1 μg) was reverse transcribed using the SuperScript VILO cDNA Synthesis kit (Invitrogen). The synthesized cDNA was used for quantitative polymerase chain reaction (qPCR) on a 7900HT Real-Time cycler (Applied Biosystem) as per the manufacturer’s instructions. Values were normalized using GAPDH as a housekeeping gene. Relative gene expression was calculated using the standard 2-ΔΔCT method. Primers were retrieved from the online PrimerBank database (http://pga.mgh.harvard.edu/primerbank/) as follows; Human *GAPDH*, 5’- GGAGCGAGATCCCTCCAAAAT-3’ and 5’- GGCTGTTGTCATACTTCTCATGG-3’; Human *CCR2*, 5’-TACGGTGCTCCCTGTCATAAA-3’ and 5’-TAAGATGAGGACGACCAGCAT-3’; Human *CX3CR1*, 5’-ACTTTGAGTACGATGATTTGGCT-3’ and 5’-GGTAAATGTCGGTGACAC TCTT-3’; Mouse *Gapdh*, 5’-AGAACATCATCCCTGCATCC-3’ and 5’-CACATTGGGGGTAGGA ACAC-3’; Mouse *Ccr2*, 5’-ATCCACGGCATACTATCAACATC-3’ and 5’-TCGTAGTCATACGG TGTGGTG-3’; Mouse *Cx3cr1*, 5’-GAGTATGACGATTCTGCTGAGG-3’ and 5’-CAGACCGAAC GTGAAGACGAG-3’.

### Flow Cytometry and Sorting

Total blood or bone marrow was incubated with 1x red blood cell (RBC) Lysis Buffer (Biolegend) for 10 min on ice. Cells were washed and re-suspended in PBS including 1% (w/v) bovine serum albumin (BSA, Sigma-Aldrich). 5×10^5^ cells (or 2×10^6^ cells for human bone marrow cells) in 100 μL of PBS including 1% (w/v) BSA were incubated with 10% (v/v) human serum (Sigma Aldrich; for human cells) or 1% (v/v) CD16/CD32 Fc blocking reagent (BD Pharmingen; for mouse cells) for 1 hour on ice to block Fc receptors. Cells were stained with fluorophore-conjugated antibodies for 30 min on ice. For human cells, the following antibodies were used: CD3-BV711 (OKT3), CD56-BV711 (HCD56), CD19-BV711 (HIB19), CD14-BV510 (M5E2), HLA-DR-BV650 (L243), CX3CR1-FITC (2A9-1), CCR2-PE-Cy7 (KO36c2), CCR1-PE (5F10B29) from Biolegend, CD45-PE-Texas Red (HI30) from Invitrogen, and CD16-EF450 (EBioCB16) from eBioscience. To validate bone marrow purity, CD34-APC (581) from Biolegend, CD71-VioBlue (AC102) from Miltenyi, and CD235a-FITC (JC159) from Acris were used. For mouse cells, the following antibodies were used: CD45-PE Dazzle (30-F11), CD11b-BV650 (M1/70), CD115-BV605 (AFS98), Ly6C-Pacific Blue (HK1.4), Ly6G-BV510 (1A8), CX3CR1-PECy7, -FITC and -APC (SA011F11) from Biolegend, CCR1-PE (643854), CCR2-PE (475301) and CX3CR1-PE (FAB585P) from R&D Systems. For intracellular staining, cells stained with antibodies for surface markers were incubated with Fixation Permeabilization Reagent (eBioscience) for 30 min at room temperature, and then re-suspended in 1x Permeabilization buffer (eBioscience). The permeabilized cells were blocked with 2% (v/v) normal rat serum, and stained with anti-CX3CR1 antibodies for 30 min at room temperature. To detect the binding of fluorescent ligand, 5×10^5^ of human or mouse blood cells were incubated in αMEM (Gibco) including 10% (v/v) FBS with or without 250ng/mL of APC-conjugated recombinant human CX3CL1 (Almac) at 4 °C for 1 hour. The incubated cells were re-suspended in PBS including 1% (v/v) BSA, and stained with antibodies for lineage markers as described above. At least 10,000 events in the monocyte gate were acquired by LSR Fortessa (BD Biosciences), and results were analyzed using FlowJo software (FlowJo v10.X; TreeStar) or DIVA software (BD Biosciences). Cell sorting was performed at 4°C in 1.5 ml RNAse and DNAse free tubes (Simport) pre-filled with 750 µl of PBS including 0.1% (w/v) BSA. A purity of sorted cells was checked every time. A minimum of 5,000 events in the monocyte/macrophage gate was acquired for cytofluorimetric analysis. Results were analyzed with FlowJo (Treestar) or DIVA software (BD). All antibodies were titrated before use.

### Microarray Analysis

Human and mouse monocyte microarray data (GSE18565 and GSE17256 respectively) were downloaded from GEO using GEO2R. All statistical calculations were performed in R language (version 3.5). Human and mouse samples were processed individually. Samples were normalized using the limma function normalizeBetweenArrays [12]. Probes representing the same gene were averaged to a single value. Differential expression analysis was performed using the limma package and significantly differentially expressed genes were selected with controlled False Positive Rate (B&H method) at 5% (FDR ≤ 0.05). Genes with log2FC ≥ +0.5 or log2FC ≤ −0.5 were considered to be up-regulated or down-regulated in C-Mo compared to NC-Mo respectively.

### Statistics

Statistical significance was calculated by Student’s *t*-test for two group comparison or by one-way or two-way ANOVA when comparing three or more groups. *P*-values ≤ 0.05 were considered as statistically significant.

## RESULTS

### Expression of *Cx3cr1* mRNA is not Different between Classical and Non-classical Monocytes in Mouse Blood

We first performed a side-by-side comparison in *CCR2* and *CX3CR1* mRNA expression between human and mouse monocyte subsets in the blood. In human samples, we selected monocytes characterized as CD45^+^CD3^−^CD19^−^CD56^−^HLA-DR^+^ and subdivided them into CD14^+^CD16^−^ (C-Mo) and CD14^dim^CD16^+^ (NC-Mo) populations. In mouse samples, monocytes characterized as CD45^+^CD115^+^Ly6G^−^ were subdivided into CD11b^+^Ly6C^high^ (C-Mo) and CD11b^+^Ly6C^low^ (NC-Mo) populations (**Supplementary Figure 2**). Consistent with previous reports [8], we found that C-Mo expressed higher levels of *CCR2* mRNA than NC-Mo both in human and mouse (**Figure 1A**). We also measured *CX3CR1* expression in NC-Mo compared to C-Mo in human. However, despite obvious differences in expression in human NC-Mos compared to C-Mos there was no difference in mice difference in *Cx3cr1* mRNA levels between the monocyte subsets in mice (**Figure 1B**). We further analyzed a published microarray data set in which gene expression profiles between human and mouse monocyte subsets were compared side-by-side [13]. Consistent with our qPCR data, differential expression of *CCR2* (C-Mo > NC-Mo) was found both in human and mouse whereas significant differential *CX3CR1* expression (C-Mo < NC-Mo) was found only in humans (**Supplementary Figure 3**); murine CX3CR1 expression was detected in both C-Mo and NC-Mo but the difference in expression between the two subsets did not reach statistical significance (data not shown).

**Figure 1:**
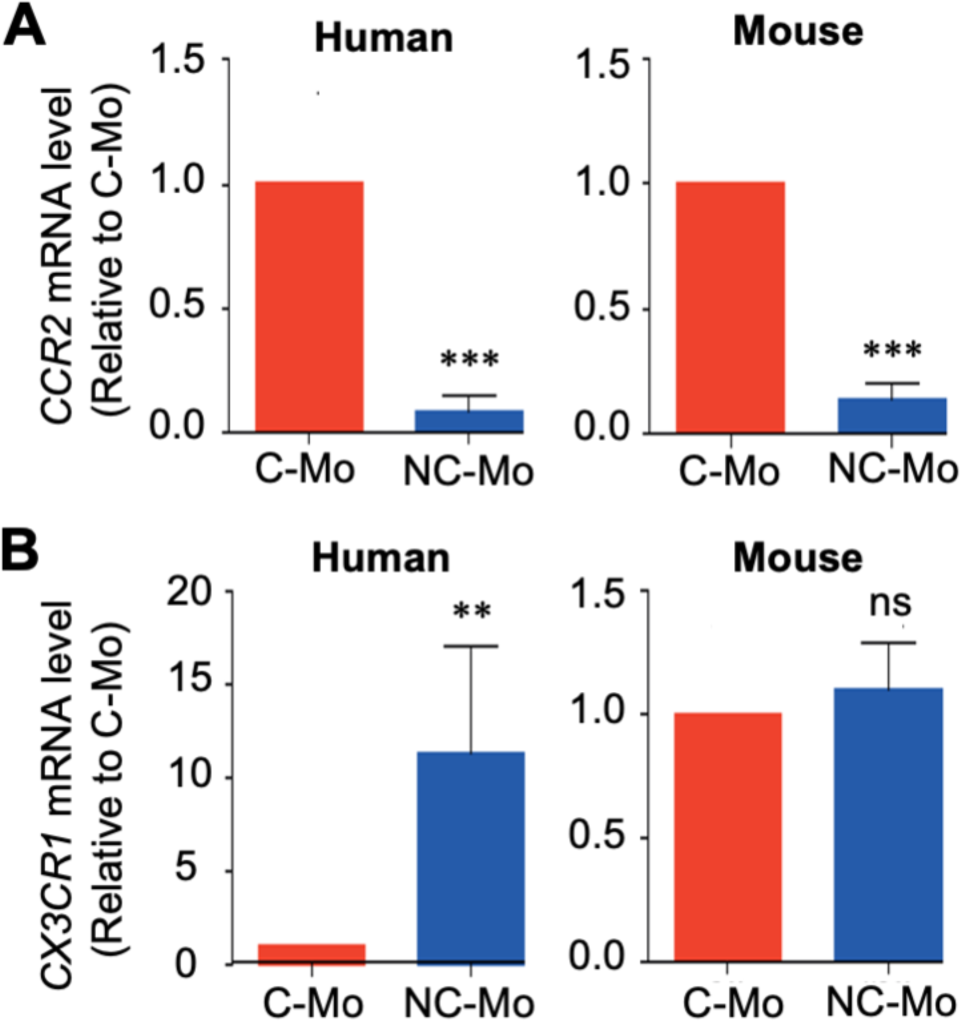
Comparison of chemokine receptor mRNA expression between monocyte subsets in human and mouse blood. **(A, B)** Expression of *CCR2* (A) and *CX3CR1* (B) mRNA in classical monocytes (C-Mo) and non-classical monocytes (NC-Mo) isolated from human and mouse blood. Data are normalized to GAPDH mRNA levels and expressed as relative to C-Mo (n =3). Data depicted as Mean±SEM. **p < 0.001 and ***p < 0.0001, Student’s t-test; ns, not significant.

### Mouse C-Mo and NC-Mo in The Blood Express Comparable Levels of CX3CR1 Protein

To further validate these findings, flow cytometric analysis of human and mouse circulating monocytes was performed to assess the protein levels of CCR2 and CX3CR1. As expected, both human and mouse C-Mo showed higher CCR2 expression compared to NC-Mo (**Figure 2A,B**). We also found that human NC-Mo expressed significantly higher levels of CX3CR1 protein than C-Mo (**Figure 2C,D**). In contrast, mouse C-Mo and NC-Mo expressed CX3CR1 protein at similar levels (**Figure 2C,D**), consistent with the mRNA expression profile. We found essentially the same results in flow cytometry using CX3CR1 monoclonal antibodies labeled with different fluorophores (PE/Cy7, APC, and FITC) or another polyclonal anti-CX3CR1 antibody (**Supplementary Figure 4**). We also found that CX3CR1 levels in monocytes detected by these antibodies were higher than those in neutrophils (**Supplementary Figure 4**), which suggest that our flow cytometry data are not biased by differences in fluorophores or quality of antibodies. We further determined intracellular levels of CX3CR1 by flow cytometry, but there were no significant differences in expression between C-Mo and NC-Mo (**Figure 2E,F**). These results indicate that neither cell surface nor intracellular expression of CX3CR1 protein is different between mouse C-Mo and NC-Mo in the blood.

**Figure 2:**
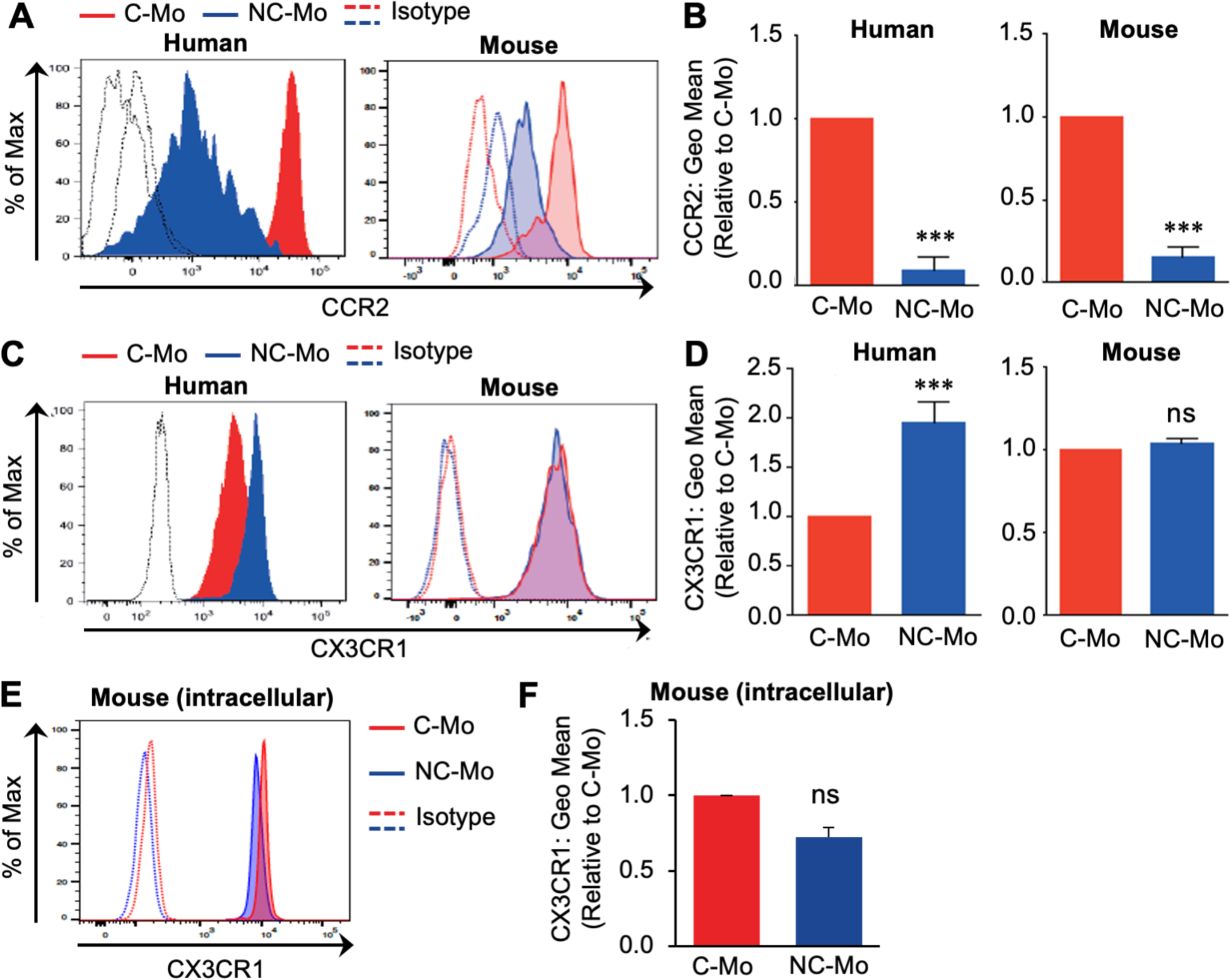
Expression of CCR2 and CX3CR1 protein on monocyte subsets in human and mouse blood. **(A)** Representative histogram showing CCR2 protein expression on human and mouse classical monocytes (C-Mo) and non-classical monocytes (NC-Mo) in the blood. Red and blue histograms indicate C-Mo and NC-Mo respectively, and dotted lines indicate their isotype control staining. **(B)** Levels of CCR2 protein (geometric mean fluorescence intensity) in human and mouse monocyte subsets in the blood. Data are expressed as fold changes relative to C-Mo (n = 5). Data depicted as Mean±SEM; ***p < 0.0001 and ****p < 0.00001, Student’s t-test. **(C)** Representative histogram showing CX3CR1 protein expression on monocyte subsets in human and mouse blood. **(D)** Levels of CX3CR1 protein on human and mouse monocyte subsets in the blood. Data are expressed as fold changes relative to C-Mo (n = 5). Data depicted as Mean±SEM; ***p < 0.0001 and ****p < 0.00001, Student’s t-test; ns, not significant. **(E)** Representative histogram showing intracellular CX3CR1 protein expression in fixed and permeabilized monocyte subsets in human and mouse blood.

### *Cx3cr1* Promoter Activities Do Not Correlate with CX3CR1 Protein Levels in Mouse Monocyte Subsets

In order to distinguish mouse monocyte subsets, the *Cx3cr1*-GFP knock-in reporter mouse model is widely utilized [10,11]. Using this model, we further verified the expression of CX3CR1 protein in mouse monocyte subsets. In this model, the GFP gene replaces the *Cx3cr1* gene. Therefore homozygous (*Cx3cr1*^GFP/GFP^) mice lack a functional *Cx3cr1* allele, which results in the reduction of NC-Mo population in the blood [10]. Consistent with previous reports, the ratio of NC-Mo population in CD45^+^ leukocytes was significantly lower in *Cx3cr1*^GFP/GFP^ mice (0.4 ± 0.2%) compared to control *Cx3cr1*^+/+^ mice (2 ± 0.3%), while it was significantly expanded in *Cx3cr1*^GFP/+^ mice (3.8 ± 0.4%; **Figure 3A**). We also confirmed that NC-Mo in *Cx3cr1*^GFP/GFP^ and *Cx3cr1*^GFP/+^ but not in wild type (*Cx3cr1*^+/+^) mice expressed high levels of GFP (**Figure 3B**). In the same samples, CX3CR1 protein expression detected by antibodies was reduced in *Cx3cr1*^GFP/+^ NC-Mo compared to *Cx3cr1*^+/+^ NC-Mo, and was almost negative in *Cx3cr1*^GFP/GFP^ NC-Mo (**Figure 3C**). Likewise, the binding of fluorescently labeled ligand (APC-conjugated recombinant human CX3CL1) on NC-Mo was decreased in *Cx3cr1*^GFP/+^ and *Cx3cr1*^GFP/GFP^ mice in accordance with the deletion of functional *Cx3cr1* gene (**Figure 3D**). These results indicate the accuracy of CX3CR1 levels detected by monoclonal antibodies or fluorescent ligands. We then compared *Cx3cr1* promoter activities and CX3CR1 protein levels between C-Mo and NC-Mo in the blood of *Cx3cr1*^+/+^, *Cx3cr1*^GFP/+^, and *Cx3cr1*^GFP/GFP^ mice. As reported [10], expression of GFP (i.e., *Cx3cr1* promoter activity) was higher in NC-Mo compared to C-Mo in *Cx3cr1*^GFP/+^ mice (**Figure 3E**; *Cx3cr1*-GFP). In contrast, there were no differences in CX3CR1 levels detected by antibodies between C-Mo and NC-Mo in all genotypes (**Figure 3E**; CX3CR1-mAb). Cell surface CX3CR1 expression indicated by binding of its fluorescent ligand CX3CL1 was also comparable between NC-Mo and C-Mo in mice (**Figure 3E**; APC-CX3CL1) whereas it was significantly higher in NC-Mo than C-Mo from human blood (**Supplementary Figure 5**). These results indicate that *Cx3cr1* promoter activities are higher in mouse NC-Mo in the blood compared to C-Mo, but that this does not correlate with mRNA or protein levels.

**Figure 3:**
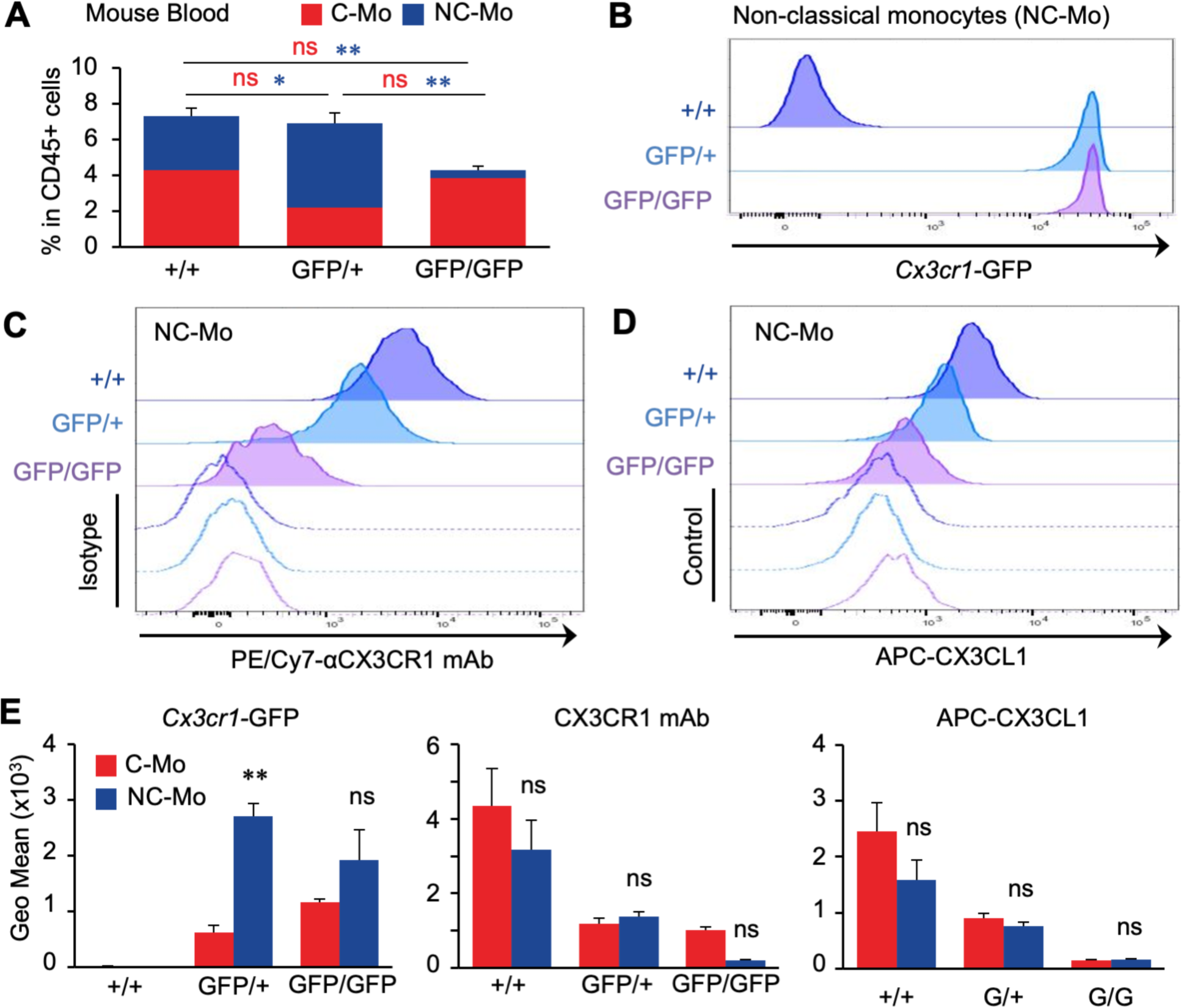
CX3CR1 expression in circulating monocyte subsets in wild type and *Cx3cr1*-GFP reporter knock-in mice. **(A)** Distribution of classical monocytes (C-Mo; red) and non-classical monocytes (NC-Mo; blue) in the blood of C57BL/6J (+/+), heterozygous knock-in *Cx3cr1*^GFP/+^ (GFP/+), and homozygous knock-in *Cx3cr1*^GFP/GFP^ (GFP/GFP) mice. Data are expressed as percentage of monocyte subsets in CD45^+^ cells (n = 3). Data depicted as Mean±SEM. *p < 0.05,**p < 0.005, Student’s t test; ns, not significant; Blue and red asterisks/ns refer to statistical test between NC- and C-Mo subsets respectively. **(B, C, D)** Representative histograms showing expression of GFP (B) or CX3CR1 that are detected by the binding of monoclonal antibodies (C; PE/Cy7-aCX3CR1 mAb) or fluorescently labeled ligand (D; APC-CX3CL1) in non-classical monocytes in the blood of C57BL/6J (+/+), *Cx3cr1*^GFP/+^ (GFP/+), and *Cx3cr1*^GFP/GFP^ (GFP/GFP) mice. Dotted lines indicate the control cells incubated with isotype control IgG (C) or PBS (D). **(E)** CX3CR1 expression in C-Mo (red) and NC-Mo (blue) detected by *Cx3cr1* promoter-driven GFP, anti-mouse CX3CR1 antibodies, and APC-CX3CL1 binding. Monocyte subsets were gated from leukocytes in the blood of C57BL/6J (+/+), *Cx3cr1*^GFP/+^ (GFP/+), and *Cx3cr1*^GFP/GFP^ (GFP/GFP) mice. Data are expressed as fold changes relative to C-Mo (n = 3). Data depicted as Mean±SEM. **p < 0.01 versus C-Mo, Student’s t-test; ns, not significant.

### Mouse NC-Mo in The Bone Marrow Express Higher Levels of CX3CR1 Protein Compared to C-Mo

It has been reported that monocytes in the bone marrow and blood have different chemokine receptor profiles. For example, a distinct population of mouse Ly6C^high^ monocytes in the bone marrow expresses high levels of CXCR4 whereas Ly6C^high^ monocytes in the blood do not express this receptor [14,15]. We therefore investigated CX3CR1 expression in monocyte subsets in the bone marrow using the *Cx3cr1*-GFP knock-in reporter mouse model (**Supplementary Figure 6**). In contrast to circulating monocytes, the Ly6C^low^ NC-Mo population was very small in the bone marrow and the ratio of NC-Mo in CD45^+^ leukocytes was comparable between *Cx3cr1*^+/+^ (0.1 ± 0.05%) and *Cx3cr1*^GFP/GFP^ mice (0.1 ± 0.05%), whereas it was significanlty increased in *Cx3cr1*^GFP/+^ mice (0.5 ± 0.07%) (**Figure 4A**). We also found that the Ly6C^high^ population was significantly expanded in *Cx3cr1*^GFP/+^ and *Cx3cr1*^GFP/GFP^ mice. Using these samples, we investigated *Cx3cr1* promoter activities and found that GFP expression was higher in NC-Mo compared to C-Mo in *Cx3cr1*^GFP/+^ mice (**Figure 4B**). We also measured CX3CR1 expression on the two subsets in the bone marrow using the monoclonal antibodies that had been used for the analyses of blood samples. Interestingly, CX3CR1 protein levels were significantly higher in NC-Mo compared to C-Mo in *Cx3cr1*^+/+^ and *Cx3cr1*^GFP/+^ mice although CX3CR1 expression in monocytes was very low in *Cx3cr1*^GFP/GFP^ mice (**Figure 4C,D**). Consistent with the mouse data, we found that human NC-Mo in the bone marrow also expressed higher levels of CX3CR1 protein compared to C-Mo (**Figure 4E,F**).

**Figure 4:**
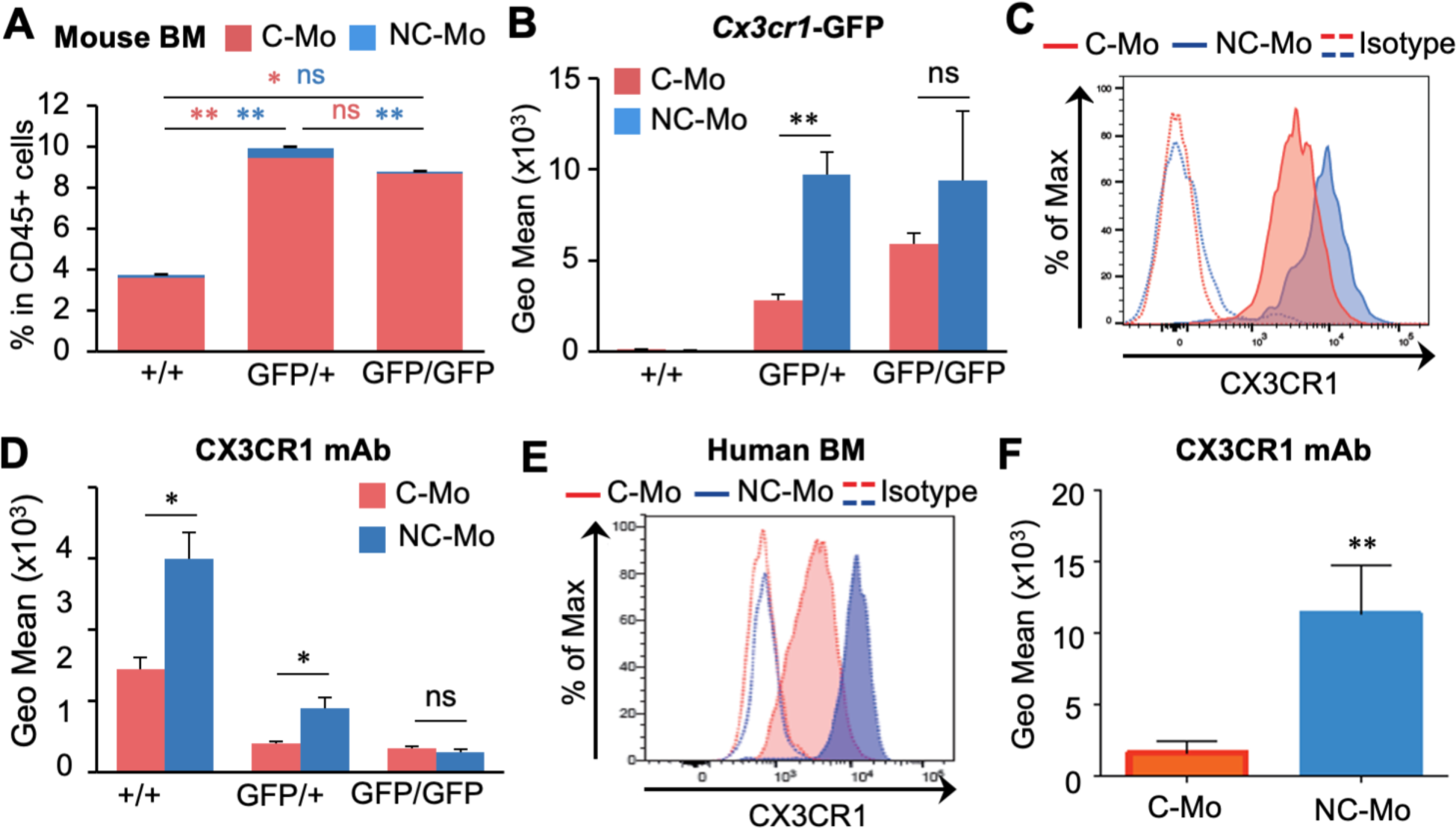
CX3CR1 expression on monocyte subsets in the bone marrow. **(A)** Distribution of classical monocytes (C-Mo; red) and non-classical monocytes (NC-Mo; blue) in the bone marrow of C57BL/6J (+/+), heterozygous knock-in *Cx3cr1*^GFP/+^ (GFP/+), and homozygous knock-in *Cx3cr1*^GFP/GFP^ (GFP/GFP) mice. Data are expressed as percentage of monocyte subsets in CD45^+^ cells (n = 3). Data depicted as Mean±SEM. *p < 0.005, **p < 0.005, Student’s t test; ns, not significant. Blue and red asterisks/ns refer to statistical test between NC- and C-Mo subsets respectively **(B)** Expression of *Cx3cr1* promoter-driven GFP in monocyte subsets in the bone marrow of C57BL/6J (+/+), *Cx3cr1*^GFP/+^ (GFP/+), and *Cx3cr1*^GFP/GFP^ (GFP/GFP) mice. Data are expressed as fold changes relative to C-Mo (n = 3). Data depicted as Mean±SEM. **p < 0.01 versus C-Mo, Student’s t-test; ns, not significant. **(C)** Representative histogram showing CX3CR1 expression detected by anti-mouse CX3CR1 monoclonal antibodies in monocyte subsets in the bone marrow of C57BL/6J mouse. **(D)** CX3CR1 levels detected by anti-mouse CX3CR1 antibodies in monocyte subsets in the bone marrow of C57BL/6J (+/+), *Cx3cr1*^GFP/+^ (GFP/+), and *Cx3cr1*^GFP/GFP^ (GFP/GFP) mice. Data are expressed as fold changes relative to C-Mo (n = 3). Data depicted as Mean±SEM. *p < 0.05 versus C-Mo, Student’s t-test; ns, not significant. **(E**) Representative histogram showing CX3CR1 expression in human C-Mo and NC-Mo in the bone marrow. **(F)** CX3CR1 levels (GEO mean) detected by anti-human CX3CR1 antibodies in human monocyte subsets in the bone marrow. Data depicted as Mean±SEM. **p < 0.0001, Student’s t test.

## DISCUSSION

Although it has been demonstrated that human and mouse genomes are very similar [16,17], leukocytes including monocytes show significant differences between human and mouse in their phenotypic marker expression, numbers, and responses to stimuli [18-20]. For example, human monocyte subpopulations C-Mo and NC-Mo are characterized by expression level of CD14 and CD16, whereas these populations are defined in mouse by the differential expression of Ly6C [8]. In mouse, NC-Mo that are believed to originate from C-Mo by monocyte conversion account for ∼50% of total monocytes in the blood [21,22], while NC-Mo in human represent only ∼10% of total monocytes [23,24]. Despite these differences, only a few reports have looked at phenotypical and/or functional comparison between human and mouse monocyte subsets [13,25]. In addition, many studies utilize mouse monocytes from the bone marrow instead of blood as a model of human peripheral blood monocytes, a situation that is not ideal for comparative immunology. In this study, we compared mouse and human monocyte subsets in the blood and bone marrow side-by-side, particularly focusing on chemokine receptor CX3CR1 that is considered as a functional receptor characteristic for NC-Mo both in human and mouse.

In contrast to what is/has been claimed by the literature, we have shown that CX3CR1 is not differentially expressed in mouse C-Mo and NC-Mo in blood, while the two subsets in the bone marrow differentially express the receptor. This discrepancy could be explained by the fact that the majority of mouse studies have used the *Cx3cr1*^GFP/+^ knock-in model and correlated the expression of GFP with the expression of CX3CR1 on the surface of monocytes [26-29]. Indeed, we confirmed that *Cx3cr1* promoter-dependent GFP expression was significantly different between mouse blood C-Mo and NC-Mo as reported previously [10]. However, the GFP level did not match the actual CX3CR1 expression on the surface of the monocytic subsets determined either by anti-CX3CR1 antibodies or by fluorescently labeled CX3CL1 ligand binding. These results indicate that, although the CX3CR1 promoter activity is different in C-Mo and NC-Mo as reported by GFP expression, the CX3CR1 protein production is equalized in both monocytic subsets due to different mRNA stability or post-translational mechanisms.

Validation at CX3CR1 protein level on blood monocytes also demonstrated that CX3CR1 expression profile in human monocytes is different from that in mouse monocytes, i.e., human NC-Mo showed higher expression of CX3CR1 protein compared to C-Mo. Using a microarray approach, Ingersoll *et al*. have profiled transcriptional landscapes of human and mouse blood monocytes [13]. In this unique microarray dataset, *CX3CR1* mRNA expression is significantly higher in NC-Mo than C-Mo in human but not in mouse (**Supplementary Figure 3**), which is consistent with our data. These findings suggest that a regulatory mechanism behind CX3CR1 expression is different between human and mouse monocytes.

In conclusion this study demonstrated that C-Mo and NC-Mo in mouse blood do not differentially express CX3CR1 unlike those in human blood. Our data gives a “note of caution” for future nomenclature of mouse monocyte subsets, as these cells cannot be defined as CX3CR1^high^ or CX3XR1^low^ based only on GFP expression in the *Cx3cr1*^GFP/+^ knock-in model. Our results also suggest that consideration should be given when studying the role of monocyte subsets in human diseases using mouse models. CX3CR1 signaling is an essential survival factor for NC-Mo but not C-Mo as demonstrated by several studies [26,30,31]. Further studies are now needed to compare the intracellular signaling activity of CX3CR1 in mouse and human C-Mo and NC-Mo.

## ABBREVIATIONS

CCR2: CC-chemokine receptor 2
C-Mo: classical monocytes
CX3CR1: CX3C-chemokine receptor 1
GFP: green fluorescent protein
NC-Mo: non-classical monocytes
qPCR: quantitative polymerase chain reaction

## AUTHOR CONTRIBUTIONS

LC, TK, SF, EH, NK, GM performed the experiments and data analysis. LC, TK, EH and JWP wrote the manuscript. All authors critically revised the manuscript. The authors declare no competing financial interests.

## ACKNOWLEDGMENTS

This research was supported by Wellcome Trust (101067/Z/13/Z) and MRC Centre grant MR/N022556/1 to JWP. Flow cytometry data was generated with support from the QMRI Flow Cytometry and Cell Sorting Facility (University of Edinburgh, UK) and the Central Facility of Sanquin (Sanquin Research, the Netherlands). We would like to thank the CIR blood resource (AMREC #15-HV-013) for the recruitment of blood from normal controls. We also thank Dr. Bain for useful scientific discussions.

## CONFLICT OF INTERESTS

The authors declare no conflict of interests.

## Supplemental Figures

**Figure S1.**
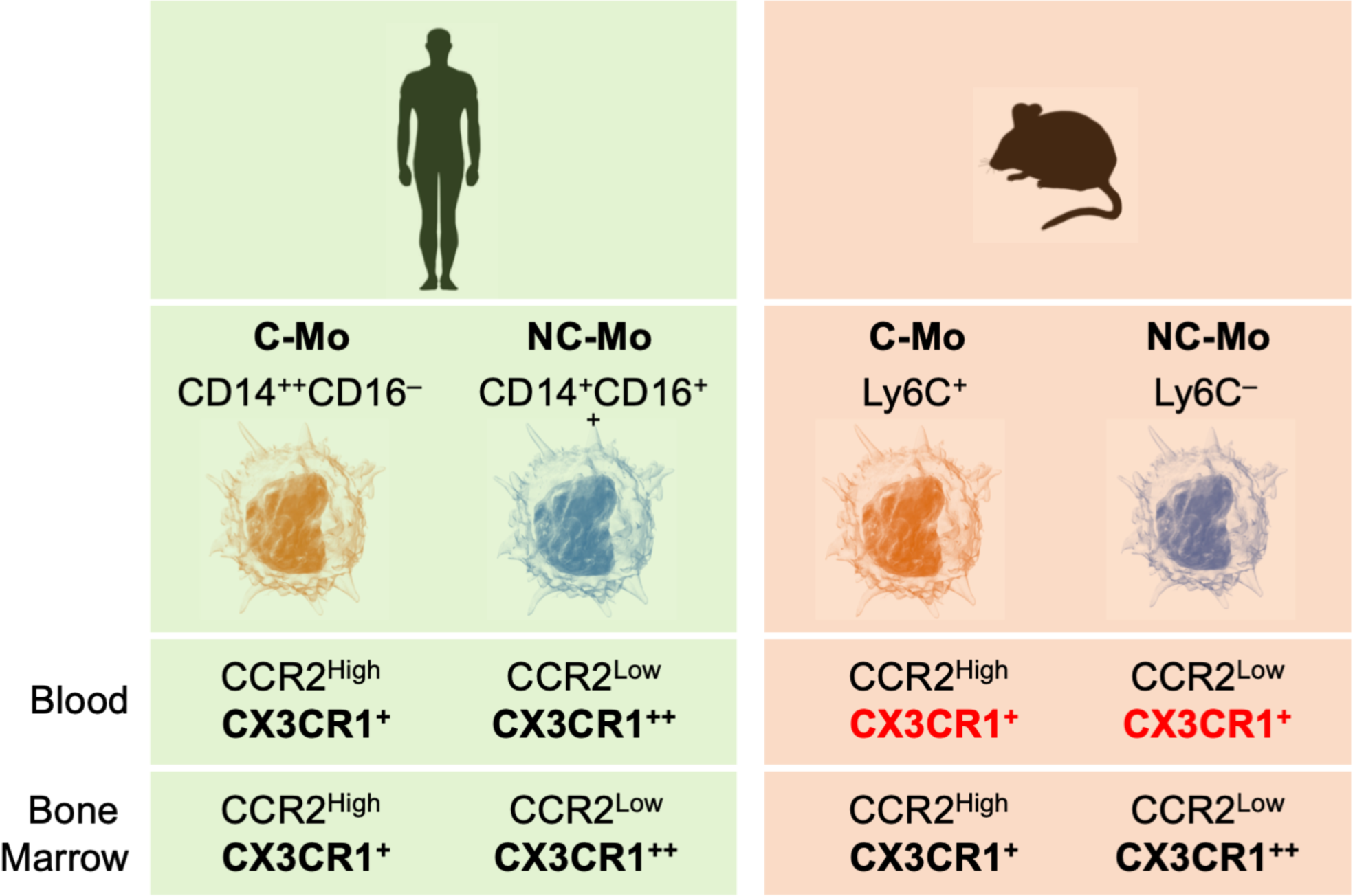
Graphical abstract.

**Figure S2.**
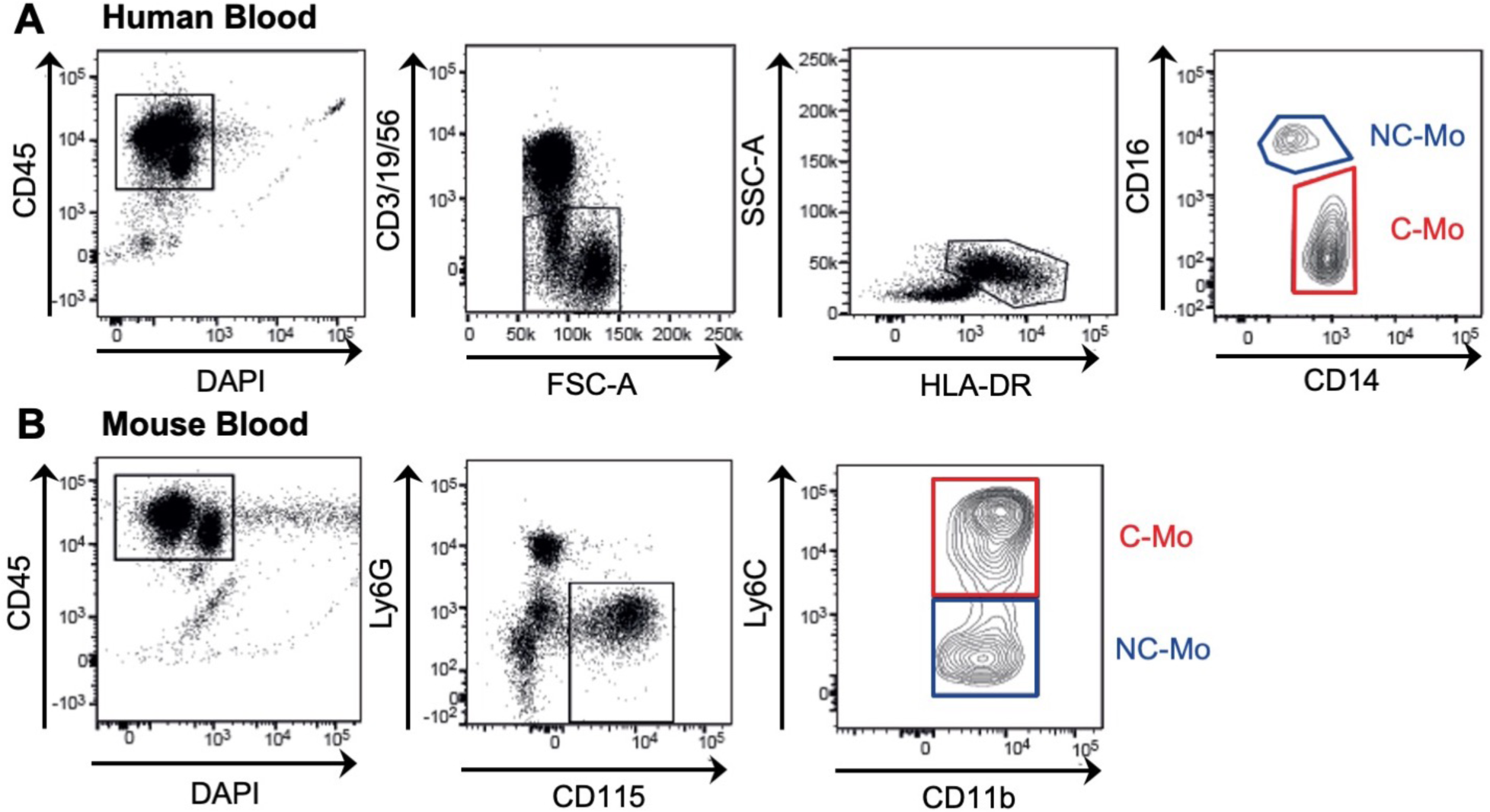
Gating strategies to identify monocyte subsets in the human and mouse blood. **(A)** Representative dot plots showing immune cells in the human blood. Monocytes were identified as CD45^+^CD3^−^CD19^−^CD56^−^HLA-DR^+^ and subdivided into classical monocytes (C-Mo; CD14^+^CD16^−^) and non-classical monocytes (NC-Mo; CD14^dim^CD16^+^). **(B)** Representative dot plots showing immune cells in the mouse blood. Monocytes were identified as CD45^+^Ly6G^−^CD115^+^ and subdivided into C-Mo (CD11b^+^Ly6C^high^) and NC-Mo (CD11b^+^Ly6C^low^).

**Figure S3.**
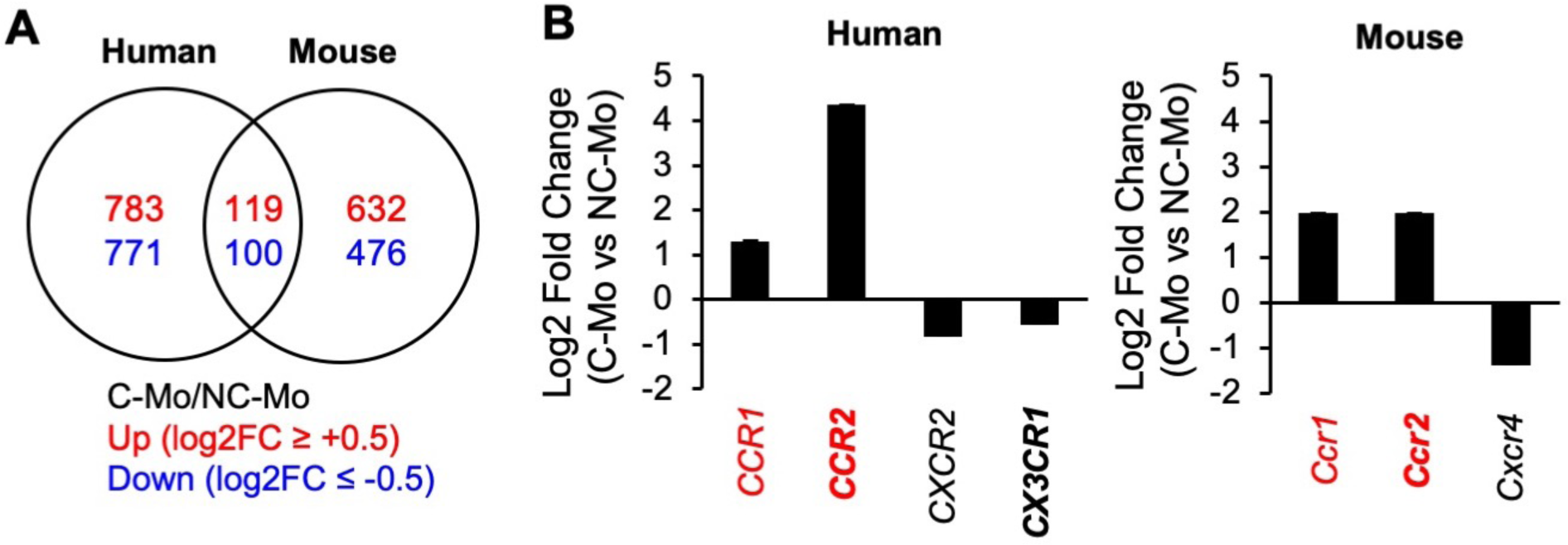
Chemokine receptor genes that are differentially expressed by human and mouse monocyte subsets in published microarray data sets. **(A)** Venn diagram of the commonly regulated genes (log2FC more or less than −1/1, FDR < 0.05) in human and mouse classical monocytes (C-Mo) compared with non-classical monocytes (NC-Mo). Red, number of up-regulated genes; blue, number of down-regulated genes. Microarray data in the GSE18565 and GSE17256 datasets were analyzed. **(B)** Genes encoding chemokine receptors which were differentially expressed by human (left) and mouse (right) C-Mo compared with NC-Mo (log2FC more or less than −1/1, FDR < 0.05). Red, genes that are commonly up-regulated in human and mouse.

**Figure S4.**
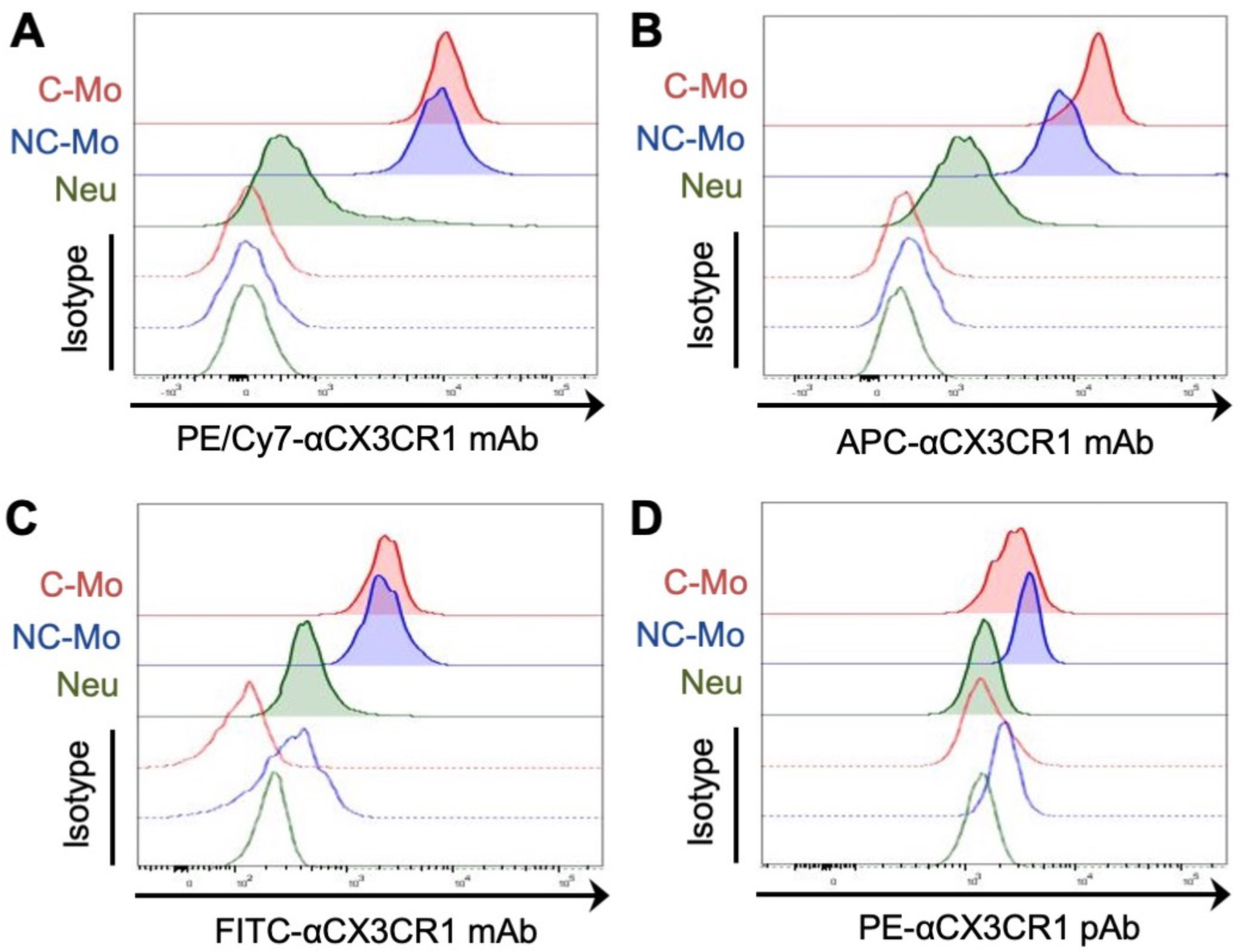
Detection of CX3CR1 by different antibodies in monocytes and neutrophils in the mouse blood. Representative histograms showing CX3CR1 protein expression in mouse classical monocytes (C-Mo), non-classical monocytes (NC-Mo), and neutrophils (Neu) detected by monoclonal antibodies conjugated with PE/Cy7 **(A)**, APC **(B)**, or FITC **(C)**, as well as polyclonal antibodies conjugated with PE as defined in material and methods **(D)**.

**Figure S5.**
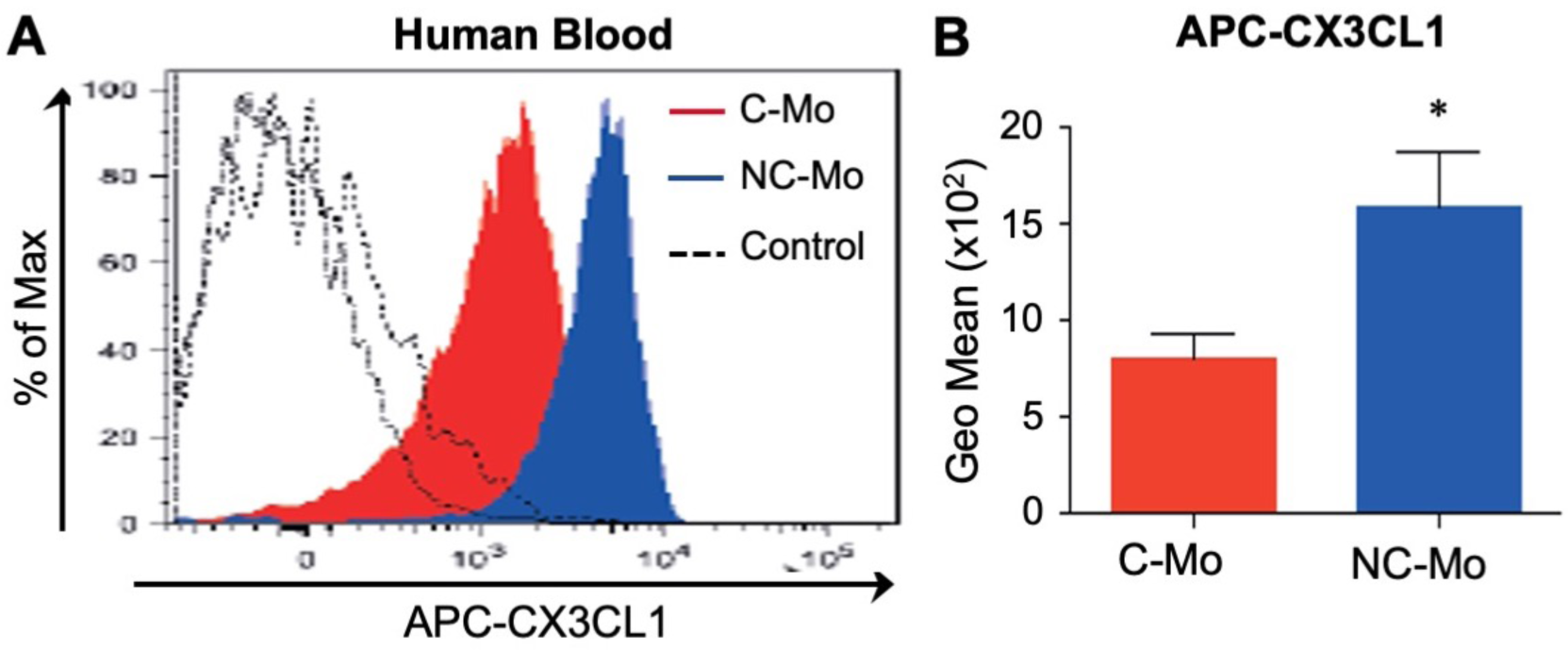
Binding of APC conjugated CX3CL1 to human monocyte subsets. **(A)** Representative histogram showing expression of CX3CR1 in classical monocytes (C-Mo) and non-classical monocytes (NC-Mo) in human blood that are detected by the binding of fluorescently labeled recombinant human CX3CL1. Dotted lines indicate the control cells incubated with PBS. **(B)** CX3CR1 levels (geometric mean fluorescence intensity) in human C-Mo (red) and NC-Mo (blue) detected by APC-CX3CL1 binding (n = 3). Data depicted as Mean±SEM. **p < 0.01 versus C-Mo.

**Figure S6.**
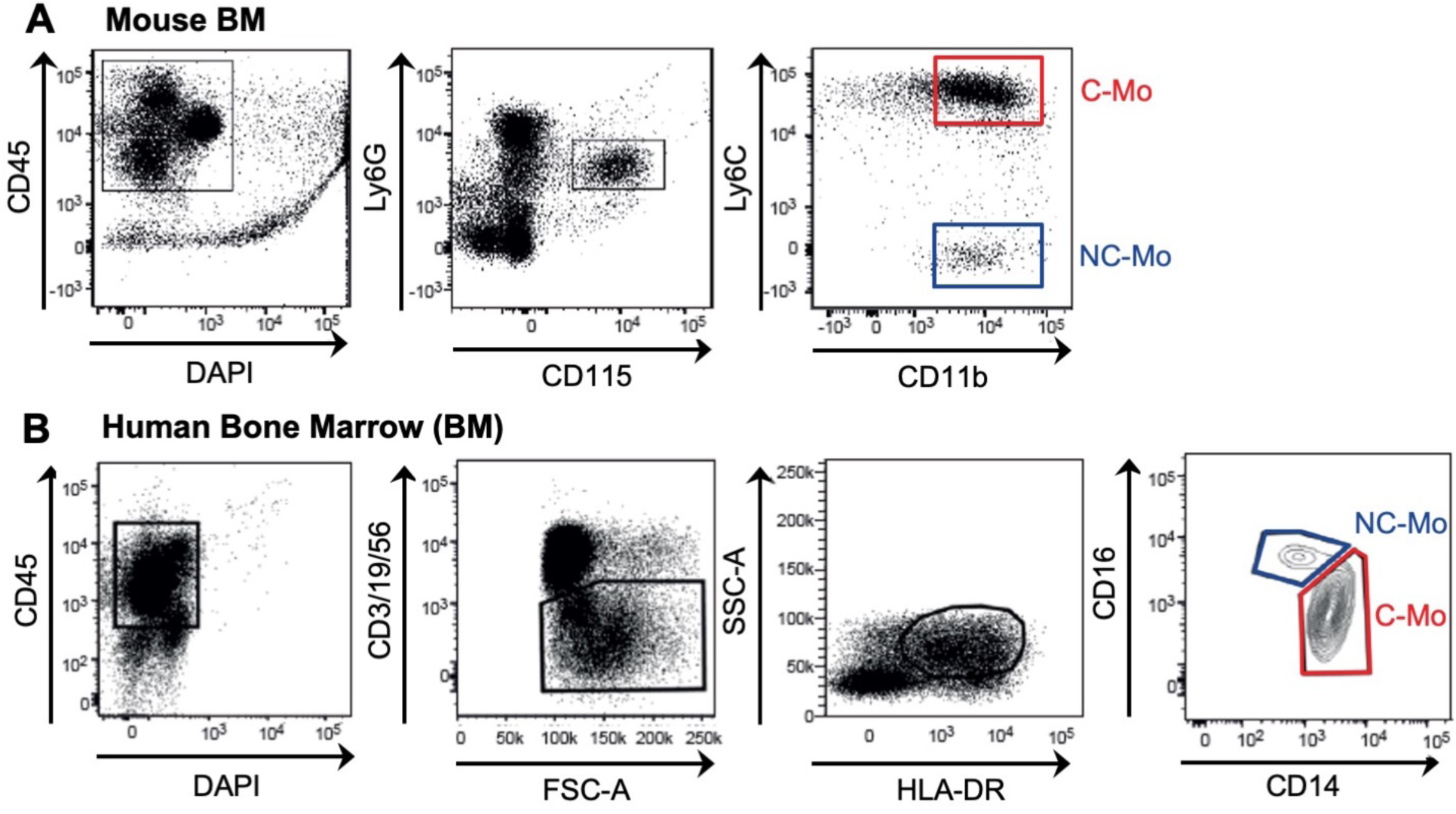
Gating strategies to identify monocyte subsets in the human and mouse bone marrow. **(A)** Representative dot plots showing immune cells in the mouse bone marrow. Monocytes were identified as CD45^+^Ly6G^−^CD115^+^ and subdivided into C-Mo (CD11b^+^Ly6C^high^) and NC-Mo (CD11b^+^Ly6C^low^). **(B)** Representative dot plots showing immune cells in the human bone marrow. Monocytes were identified as CD45^+^CD3^−^CD19^−^CD56^−^HLA-DR^+^ and subdivided into classical monocytes (C-Mo; CD14^+^CD16^−^) and non-classical monocytes (NC-Mo; CD14^dim^CD16^+^).

## Notes

### Competing Interest Statement

The authors have declared no competing interest.

### Summary of Updates

Formatting is now complete.

## REFERENCES

1. Passlick B, Flieger D, Ziegler-Heitbrock HW. Identification and characterization of a novel monocyte subpopulation in human peripheral blood. Blood (1989) 74: 2527–2534.

2. Ziegler-Heitbrock HW, Fingerle G, Ströbel M, Schraut W, Stelter F, Schütt C, et al. The novel subset of CD14+/CD16+ blood monocytes exhibits features of tissue macrophages. Eur J Immunol (1993) 23: 2053–2058. doi: 10.1002/eji.1830230902

3. Kratofil RM, Kubes P, Deniset JF. Monocyte conversion during inflammation and injury. Arterioscler Thromb Vasc Biol. (2017) 37:35–42. doi: 10.1161/ATVBAHA.116.308198.

4. Kitamura T, Qian B-Z, Pollard JW. Immune cell promotion of metastasis. Nat. Rev. Immunol. (2015) 15:73–86. doi:10.1038/nri3789.

5. Auffray C, Fogg D, Garfa M, Elain G, Join-Lambert O, Kayal S, et al. Monitoring of blood vessels and tissues by a population of monocytes with patrolling behavior. Science (2007) 317:666–70. doi: 10.1126/science.1142883

6. Cros J, Cagnard N, Woollard K, Patey N, Zhang SY, Senechal B, et al. Human CD14dim monocytes patrol and sense nucleic acids and viruses via TLR7 and TLR8 receptors. Immunity (2010) 33:375–86. doi: 10.1016/j.immuni.2010.08.012.

7. Hanna RN, Cekic C, Sag D, Tacke R, Thomas GD, Nowyhed H, et al. Patrolling monocytes control tumor metastasis to the lung. Science. (2015) 350:985–90. doi: 10.1126/science.aac9407.

8. Gordon S, Taylor PR. Monocyte and macrophage heterogeneity. Nat Rev Immunol. (2005) 5:953–964. doi: 10.1038/nri1733.

9. Weber C, Belge KU, von Hundelshausen P, Draude G, Steppich B, Mack M, et al. Differential chemokine receptor expression and function in human monocyte subpopulations. J Leukoc Biol. (2000) 67:699–704. doi: 10.1002/jlb.67.5.699.

10. Geissmann F, Jung S, Littman DR. Blood monocytes consist of two principal subsets with distinct migratory properties. Immunity. (2003) 19:71–82.

11. Jung S, Aliberti J, Graemmel P, Sunshine MJ, Kreutzberg GW, Sher A, et al. Analysis of fractalkine receptor CX(3)CR1 function by targeted deletion and green fluorescent protein reporter gene insertion. Mol Cell Biol (2000) 20:4106–4114. doi: 10.1128/mcb.20.11.4106-4114.2000.

12. Ritchie, M. E., Phipson, B., Wu, D., Hu, Y., Law, C. W., Shi, W., Smyth, G. K. (2015) limma powers differential expression analyses for RNA-sequencing and microarray studies. Nucleic Acids Res 43, e47. doi: 10.1093/nar/gkv007.

13. Ingersoll MA, Spanbroek R, Lottaz C, Gautier EL, Frankenberger M, Hoffmann R, et al. Comparison of gene expression profiles between human and mouse monocyte subsets. Blood (2010) 115: e10–19.

14. Schlueter AJ, Glasgow JK. Phenotypic comparison of multiple monocyte-related populations in murine peripheral blood and bone marrow. Cytometry A. (2006) 69:281–290. doi: 10.1002/cyto.a.20262.

15. Chong SZ, Evrard M, Devi S, Chen J, Lim JY, See P, et al. CXCR4 identifies transitional bone marrow premonocytes that replenish the mature monocyte pool for peripheral responses. J Exp Med. (2016) 213:2293–2314. doi: 10.1084/jem.20160800.

16. Waterston RH, Lindblad-Toh K, Birney E, Rogers J, Abril JF, Agarwal P, et al. Initial sequencing and comparative analysis of the mouse genome. Nature. (2002) 420:520–62. doi: 10.1038/nature01262

17. Pennacchio LA. Insights from human/mouse genome comparisons. Mamm Genome. (2003) 14:429–36. doi: 10.1007/s00335-002-4001-1.

18. Haley PJ. Species differences in the structure and function of the immune system. Toxicology (2003) 188: 49–71. doi: 10.1016/s0300-483x(03)00043-x

19. Mestas J, Hughes CC. Of mice and not men: differences between mouse and human immunology. J Immunol (2004) 172: 2731–2738.

20. Haley PJ. The lymphoid system: a review of species differences. J Toxicol Pathol (2017) 30: 111–123.

21. Sunderkötter C, Nikolic T, Dillon MJ, Van Rooijen N, Stehling M, Drevets DA, et al. Subpopulations of mouse blood monocytes differ in maturation stage and inflammatory response. J Immunol. (2004) 172:4410–4407. doi: 10.4049/jimmunol.172.7.4410

22. Yona S, Kim KW, Wolf Y, Mildner A, Varol D, Breker M, et al. Fate mapping reveals origins and dynamics of monocytes and tissue macrophages under homeostasis. Immunity (2013) 38: 79–91. doi: 10.1016/j.immuni.2012.12.001.

23. Mukherjee R, Kanti Barman P, Kumar Thatoi P, Tripathy R, Kumar Das B, Ravindran B. Non-Classical monocytes display inflammatory features: validation in sepsis and systemic lupus erythematous. Sci Rep. (2015) 5:13886. doi: 10.1038/srep13886.

24. Patel AA, Zhang Y, Fullerton JN, Boelen L, Rongvaux A, Maini AA, et al. The fate and lifespan of human monocyte subsets in steady state and systemic inflammation. J Exp Med. (2017) 214:1913–1923. doi: 10.1084/jem.20170355.

25. Reynolds G, Haniffa M. Human and mouse mononuclear phagocyte networks: a tale of two species? Front Immunol (2015) 6:330.

26. Landsman L, Bar-On L, Zernecke A, Kim KW, Krauthgamer R, Shagdarsuren E, et al. CX3CR1 is required for monocyte homeostasis and atherogenesis by promoting cell survival. Blood. (2009) 113:963–972. doi: 10.1182/blood-2008-07-170787.

27. Medina-Contreras O, Geem D, Laur O, Williams IR, Lira SA, Nusrat A, et al. CX3CR1 regulates intestinal macrophage homeostasis, bacterial translocation, and colitogenic Th17 responses in mice. J Clin Invest. (2011) 121:4787–95. doi: 10.1172/JCI59150.

28. Jacquelin S, Licata F, Dorgham K, Hermand P, Poupel L, Guyon E, et al. CX3CR1 reduces Ly6Chigh-monocyte motility within and release from the bone marrow after chemotherapy in mice. Blood. (2013) 122:674–683. doi: 10.1182/blood-2013-01-480749.

29. McArdle S, Mikulski Z, Ley K. Live cell imaging to understand monocyte, macrophage, and dendritic cell function in atherosclerosis. J Exp Med. (2016) 213:1117–1131. doi: 10.1084/jem.20151885.

30. Feng X, Szulzewsky F, Yerevanian A, Chen Z, Heinzmann D, Rasmussen RD, et al. Loss of CX3CR1 increases accumulation of inflammatory monocytes and promotes gliomagenesis. Oncotarget (2015) 6:15077–15094. doi: 10.18632/oncotarget.3730

31. Collar AL, Swamydas M, O’Hayre M, Sajib MS, Hoffman KW, Singh SP, et al. The homozygous CX3CR1-M280 mutation impairs human monocyte survival. JCI Insight (2018) 3. pii: 95417. doi: 10.1172/jci.insight.95417.

